# Exploring high-resolution cryo-ET and subtomogram averaging capabilities of contemporary DEDs

**DOI:** 10.1101/2022.01.10.475481

**Authors:** Martin Obr, Wim JH Hagen, Robert A Dick, Lingbo Yu, Abhay Kotecha, Florian KM Schur

## Abstract

The potential of energy filtering and direct electron detection for cryo-electron microscopy (cryo-EM) image processing has been well documented for single particle analysis (SPA). Here, we assess the performance of recently introduced hardware for cryo-electron tomography (cryo-ET) and subtomogram averaging (STA), an increasingly popular structural determination method for complex 3D specimens. We acquired cryo-ET datasets of EIAV virus-like particles (VLPs) on two contemporary cryo-EM systems equipped with different energy filters and direct electron detectors (DED), specifically a Krios G4, equipped with a cold field emission gun (CFEG), Thermo Fisher Scientific Selectris X energy filter, and a Falcon 4 DED; and a Krios G3i, with a Schottky field emission gun (XFEG), a Gatan Bioquantum energy filter, and a K3 DED. We performed constrained cross-correlation-based STA on equally sized datasets acquired on the respective systems. The resulting EIAV CA hexamer reconstructions show that both systems perform comparably in the 4-6 Å resolution range. In addition, by employing a recently introduced multiparticle refinement approach, we obtained a reconstruction of the EIAV CA hexamer at 2.9 Å. Our results demonstrate the potential of the new generation of energy filters and DEDs for STA, and the effects of using different processing pipelines on their STA outcomes.

## Introduction

Cryo-electron tomography (cryo-ET) is used to visualize complex biological environments in 3D. In combination with the image processing technique subtomogram averaging (STA), structures in their native context, such as the interior of cells, can be determined (Schur, 2019; Wan and Briggs, 2016). In recent years, substantial developments in cryo-ET data acquisition and STA processing have resulted in high-resolution structures of challenging samples that are not accessible by any other method (Bäuerlein and Baumeister, 2021). However, while high-resolutions at ~4Å can now be reached for some specimens, such as ribosomes (O’Reilly et al., 2020; Tegunov et al., 2021), selected virus assemblies (Dick et al., 2020; Schur et al., 2016), and apoferritin or dNTPase *in vitro* (Bouvette et al., 2021; Tegunov et al., 2021), the average resolution of cryo-ET-derived structures in the electron microscopy databank (EMDB) has only marginally increased over the last few years (with an average resolution of ~21 Å in 2021) and has remained behind what is now commonly achieved in single-particle particle (SPA) cryo-EM (~5.3 Å average resolution in 2021).

Still, the potential of cryo-ET and its increasing popularity has lead to a growing scientific community that is actively contributing to developments (for examples see (Chen et al., 2019; Himes and Zhang, 2018; Sanchez et al., 2020; Scaramuzza and Castaño-Díez, 2021; Tegunov et al., 2021)). Improving the attainable resolution and especially reducing the number of particles required to reach a given resolution via cryo-ET and STA is a matter of significant interest to the cryo-EM field.

Reasons for the overall lower resolution achieved in cryo-ET include the higher complexity specimen environments and larger, more variable macromolecular complexes, resulting in higher demands on data acquisition and image processing. Tilt series acquisition and cryo-ET data processing with the goal of obtaining high resolutions is challenging due to several limitations inherent to the method (Schur, 2019), some of these limitations are: 1) the requirement to distribute the available cumulative dose over a finite number of tilt angles, necessitating a very low exposure per image, resulting in a low signal-to-noise ratio (SNR) for each tilt; 2) increased thickness of the sample, particularly at high tilts; 3) the accumulated beam-induced damage during acquisition; 4) an incomplete sampling of the tilt range, due to mechanical limitations of the stage; and 5) lower throughput due to longer acquisition time. The first three limitations cause an increased loss of high spatial frequency information in later tilts.

Of the above-mentioned limitations, the last point has recently been addressed in several developments that exploit efficient data acquisition methods, more stable stages and faster cameras (Bouvette et al., 2021; Chreifi et al., 2019; Eisenstein et al., 2019). This has allowed obtaining datasets of hundreds of tomograms within a short time. However, points 1-4 constitute inherent limitations of cryo-ET, which have been shown to be most efficiently mitigated by better means of data acquisition. This includes using direct electron detectors (DEDs) with improved camera detective quantum efficiency (DQE) and energy-filters that remove inelastically scattered electrons, which leads to enhanced SNR. In addition, optimized acquisition schemes have been shown to mitigate these limitations to a certain degree (Bouvette et al., 2021; Hagen et al., 2017; Turoňová et al., 2020).

SPA cryo-EM has also benefited from a series of developments, such as improved electron sources, improved specimen holders, and better direct detection with higher camera frame rates. Recently, the new generation of DEDs and energy filters of two microscope hardware manufacturers (ThermoFisher Scientific (TFS) with their combination of the Selectris X energy filter and Falcon 4 DED; and Gatan with their combination of the Bioquantum energy filter and K3 DED) led to exciting results in SPA, reaching to atomic resolution for apoferritin samples (Nakane et al., 2020; Zhang et al., 2020). Another recent study also achieved similar breakthrough atomic resolution on the same sample using an aberration-corrected aplanatic Titan Krios and a Falcon 3 DED (Yip et al., 2020). These studies demonstrated the potential of SPA cryo-EM for structural biology of well-ordered and homogenous protein samples.

To assess the potential of current cryo-EM systems for high-resolution cryo-ET and STA, we tested two systems, each equipped with the newest generation of DEDs and energy filters. We acquired cryo-ET data, first on a Krios G4, equipped with a cold field emission gun (CFEG), Selectris X energy filter, and Falcon 4 DED (referred to as **System 1** herein, installed at the TFS RnD facility, Eindhoven, NL). We then acquired data on a Krios G3i, equipped with a high-brightness Schottky field emission gun (XFEG), Gatan Bioquantum energy filter and K3 DED (**System 2**, installed at EMBL Heidelberg, Germany).

The sample was Equine infectious anemia virus (EIAV) virus-like particles (VLPs), formed from a truncated variant of the main retrovirus structural protein Gag. The polyprotein Gag (consisting of the canonical domains matrix (MA), capsid (CA) and nucleocapsid (NC)) and its truncations (consisting of only individual domains or parts thereof) can be expressed recombinantly, purified, and then assembled *in vitro* to form VLPs reflecting architecture and organization of authentic virus particles (Bush and Vogt, 2014).

Retroviral VLPs are a well-suited sample to explore the limitations and the potential of cryo-ET and STA (Obr and Schur, 2019). In almost all cases, Gag-derived VLPs vary in size and curvature and lack global symmetry, making each particle a unique object. This makes them almost intractable for SPA cryo-EM approaches, and a relevant sample for demonstrating the high-resolution cryo-ET potential of the employed hardware and software. In particular, although lacking global symmetry, the locally symmetric lattice arrangement of the CA domain within Gag, with C2, C3, and C6 symmetry axes, facilitates STA processing. These local symmetries effectively reduce the angular search space (and hence computation time) and increase the available dataset size. In addition, using an *in vitro* assembly system permits preparation of VLPs in a defined concentration, allowing easier optimization of grid preparation.

For these reasons, retroviral VLPs were the first to reach resolutions better than 4 Å using cryo-ET and STA (Schur et al., 2016). They also have been frequently used for benchmarking of data acquisition (Turoňová et al., 2020), and have served as evaluation data for the development of various image processing software suites, such as novaCTF (Turoňová et al., 2017), Dynamo (Scaramuzza and Castaño-Díez, 2021), emClarity (Himes and Zhang, 2018) and M (Tegunov et al., 2021).

For our work, we used an EIAV Gag truncation construct with the capsid (CA), spacer peptide (SP) and nucleocapsid (NC) domains (referred to as CASPNC), which forms spherical and tubular VLPs *in vitro*. EIAV CASPNC VLPs were recently analyzed on a cryo-ET system equipped with the Gatan K2 DED, yielding a resolution of 3.7 Å (Dick et al., 2020). By using this sample, we were able to compare the results from the previous generation of DEDs with the results from the new DEDs.

For our evaluation, we employed two established pipelines for processing cryo-ET data. First, we evaluated the performance of both systems with equally sized datasets. For this, we used an approach as employed in previous papers studying structures of retroviral Gag assemblies. This approach is based on tomogram reconstruction with 3D CTF correction using novaCTF and the AV3-derived subtomogram averaging/alignment pipeline (Dick et al., 2020; Förster et al., 2005; Obr et al., 2021; Turoňová et al., 2020, 2017).

In addition, we used an approach which included consecutive 3D refinement in Relion (Bharat et al., 2015) and multiparticle refinement steps in the software M. This workflow previously resulted in the highest resolution STA structure to date (Tegunov et al., 2021).

Here we report that data obtained on System 1 and System 2 yield sub-4 Å reconstructions from relatively small datasets, and with consistent quality of their respective maps. Processing the full dataset obtained from System 1 via iterative refinement in Relion and multiparticle refinement in M resulted in a 2.9Å reconstruction, the highest resolution structure of retroviral capsid assemblies determined by cryo-ET and STA. Our analysis demonstrates that each system, as well as their unique data acquisition settings are compatible with high-resolution STA.

In order to support community-based software developments, the System 1 dataset presented herein is deposited to the EMPIAR database under the accession code EMPIAR-10889.

## Results & Discussion

### Data acquisition and evaluation

EIAV CASPNC VLPs were vitrified and screened on a Glacios TEM at IST Austria (see methods for details). Grids with appropriate VLP distribution and ice quality were then shipped to System 1 at TFS Eindhoven. Tilt series were acquired on a single grid in the electron event representation (EER) format using the TFS Tomography software. At the time of dataset processing, none of our STA software was compatible with native EER processing, so we summed the electron events into computational frames for subsequent processing.

We then sent the same grid to EMBL Heidelberg, where a second dataset was acquired on System 2 using SerialEM. By using the same VLP preparation on the same grid for acquisition on both systems, we aimed to diminish sample-related bias. At the same time, this also negatively affected the size of dataset 2, as the remaining number of appropriate available acquisition positions was limited.

Beyond being different in microscope and camera hardware the data acquisition on both systems differed in the selected slit width of the filter (10 eV for System 1, 20 eV for System 2). Since the K3 chip is rectangular, the field of view along the long axis of the K3 chip was kept equivalent to either axis of the square-shaped Falcon4 chip (see Supplementary Table 1 for details on the acquisition settings). This meant acquiring with a smaller pixel size on System 2, with potentially improved DQE, but also resulted in a reduced total area covered in a single tomogram for System 2.

**Supplementary Table 1:** Acquisition settings.

|  |  | System 1 | System 2 |
| --- | --- | --- | --- |
| Dataset |  | EIAV CANC Falcon 4 | EIAV CANC K3 |
| Acquisition settings | Microscope | FEI Titan Krios G4 | FEI Titan Krios G3i |
|  | Voltage (keV) | 300 | 300 |
|  | Electron source | E-CFEG | XFEG |
|  | Detector | Falcon 4 | Gatan K3 |
|  | Energy-filter | Selectris X | Gatan BioQuantum |
|  | Slit width (eV) | 10 | 20 |
|  | Super-resolution Mode | Yes | Yes |
|  | Å/pixel | 1.179 | 0.822 |
|  | Defocus range (µm) | -0.75 to 3.25 | -0.75 to 3.25 |
|  | Defocus step (µm) | 0.25 | 0.25 |
|  | Acquisition scheme | -60/60°, 3°<br>Dose-symmetric<br>Tomography 5.0 | -60/60°, 3°<br>Dose-symmetric<br>SerialEM |
|  | Total dose (electrons/ Å<sup>2</sup>) | ~143.5 | ~146 |
|  | Dose rate (electrons/ Å<sup>2</sup>/sec) | 3.43 | 38.2 |
|  | File Format | EER<br>(electron event<br>representation) | tif<br>(hardware summed raw<br>frames) |
|  | Frame number | Not applicable | 7 |
|  | Tilt series used for STA | 85 (84)* | 35 |

\* - 85 tilt series were used in the AV3 alignment, but only 84 were subjected to processing in Relion/M.

**Supplementary Table 2:** Dataset overview.

| System | Dataset | Symmetry | Particle number | (Sub-) tomogram reconstruction | STA Processing | FSC 0.143 (Å) | Fourier shells above 0.143 crit. |
| --- | --- | --- | --- | --- | --- | --- | --- |
| **1** |  |  |  |  |  |  |  |
|  | batch 1 | C6 | 20 000 | novaCTF | AV3 | 3.8 | 69/112 |
|  | batch 2 | C6 | 20 000 | novaCTF | AV3 | 3.9 | 68/112 |
|  | batch 3 | C6 | 20 000 | novaCTF | AV3 | 3.8 | 69/112 |
|  | full dataset | C6 | 62 282 | novaCTF | AV3 | 3.4 | 78/112 |
|  | full dataset | C6 | 77 659 | Warp | Relion, Multiparticle refinement | 2.9 | 136/170 |
|  | full dataset | C2 | 53 665 | Warp | Relion, Multiparticle refinement | 3.2 | 125/170 |
| **2** |  |  |  |  |  |  |  |
|  | batch 1 | C6 | 20 000 | novaCTF | AV3 | 3.9 | 66/112 |
|  | full dataset | C6 | 32 446 | Warp | Relion, Multiparticle refinement | 3.3 | 121/174 |
|  | full dataset | C2 | 18 596 | Warp | Relion, Multiparticle refinement | 3.6 | 112/174 |

**Supplementary Table 3:**
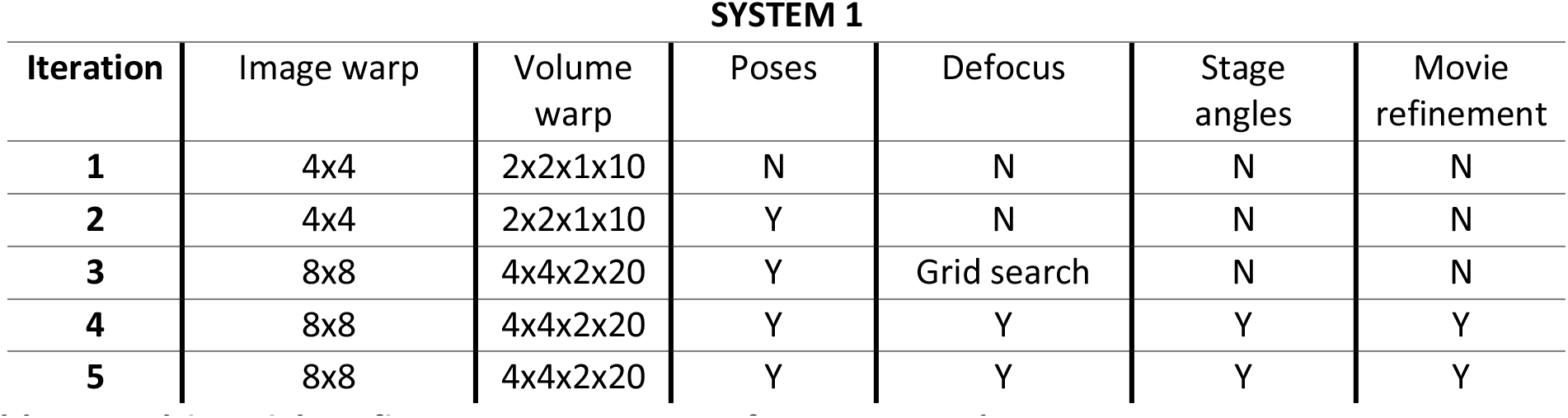
Multiparticle refinement parameters for System 1 dataset.

**Supplementary Table 4:** Multiparticle refinement parameters for System 2 dataset.

| Iteration | Image warp | Volume warp | Poses | Defocus | Stage angles | Movie refinement |
| --- | --- | --- | --- | --- | --- | --- |
| 1 | 4x3 | 2x2x1x10 | N | N | N | N |
| 2 | 4x3 | 2x2x1x10 | Y | N | N | N |
| 3 | 8x6 | 3x4x2x20 | Y | Grid search | N | N |
| 4 | 8x6 | 3x4x2x20 | Y | Y | Y | Y |
| 5 | 8x6 | 3x4x2x20 | Y | Y | Y | Y |

Tilt series in both datasets contained large amounts of spherical and tubular VLPs (Figure 1a-b), as previously reported for EIAV CASPNC assemblies (Dick et al., 2020). As expected, the hexameric arrangement of the CA domains, forming a lattice on the surface of tubes and spheres, was clearly resolved. For further evaluation of data quality, we performed defocus estimation using CTFFIND4 (Figure 1c,d). Both datasets showed accurate fitting of the contrast transfer function (CTF) up to +/- 40 degrees tilt, judged by the achieved resolution of CTF-fitting (Figure 1f,h) and the CTF fit rate (ratio between successful and all CTF fits for a given tilt angle) (Figure 1e,g). In the tilt range between −30° and +24°; and −36° and +42°, no failed CTF fits were observed in the case of System 1 and System 2, respectively (Figure 1e,g). Beyond ±40° tilt angles (the last third of the tilt series with the highest accumulated exposure dose) the successful fit rate was 67% and 86% for System 1 and System 2, respectively. Overall, data acquired on System 2 showed a slightly higher resolution of CTF fits. As there are several notable differences between the two datasets that could affect this result it is difficult to assign a reason for the slightly improved fitting results. One explanation for this could be the smaller pixel size used for the acquisition on System 2, which could have been beneficial for CTF fitting. Notably, CTF-fitting showed slightly higher reliability for defoci further from focus than 2 microns for both systems. All of the above observations suggest that both systems perform favorably at the low-dose conditions required for cryo-ET.

**Figure 1:**
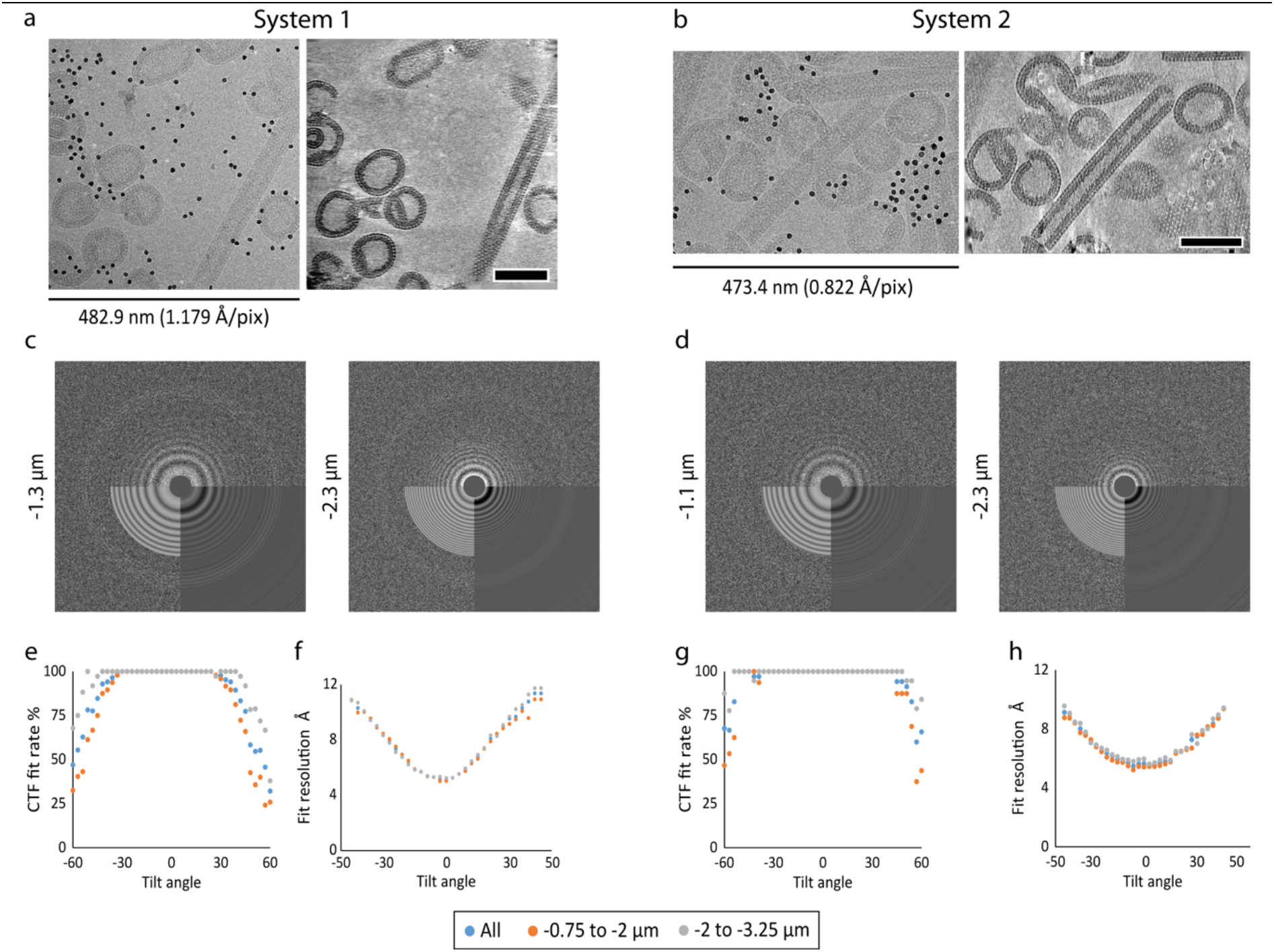
Cryo-electron tomography on current energy-filtered DEDs. **a)** Tilt image and reconstructed cryoelectron tomogram acquired on System 1. Left: 2D cryo-EM micrograph showing a single (0°) tilt. Right: 10 summed computational slices through the reconstructed tomogram. **b**) Tilt image and reconstructed cryoelectron tomogram acquired on System 2. Left: 2D cryo-EM micrograph showing a single (0°) tilt. Right: 10 summed computational slices through the reconstructed tomogram. The dimension of the x-axes is annotated for both systems. Scale bars are 100nm. **c-d)** Representative CTFFIND4 outputs for the 0° tilt of two tilt series, acquired at different defocus settings, for System 1 (c) and System 2 (d). The upper part of the power spectrum shows the experimental power spectrum, the lower right panel shows the radial average and the lower left panel shows the estimated CTF-fit. **e-h)** Success rates for CTF fitting (e,g) and the median resolution of the CTF fitting (f,h) for each tilt, described by the highest spatial frequency to which the fit was reliable. (e-f) Shows results for System 1, and (g-h) for System 2.

### STA benchmark using novaCTF

Next, to further assess the performance of the respective systems and the quality of the corresponding datasets, we performed STA on spherical VLPs. We used the subtomogram averaging workflow, as published previously (Dick et al., 2020; Turoňová et al., 2020) with minor adaptations, which utilizes 3D-CTF corrected tomograms and a constrained cross-correlation-based subtomogram averaging/alignment routine of AV3 (Förster et al., 2005).

To allow an unbiased analysis we designed a processing strategy that is not influenced by varying VLP quality. Specifically, individual spherical VLPs can differ in the completeness of their CA protein lattice (i.e. due to VLPs being broken or incompletely assembled). This results in different extents of lattice edges, which are formed by incomplete hexamers (Tan et al., 2021). This can, if present in different abundance, have a negative effect on the resolution of the final maps. In addition, the alignment of the central hexamer depends also on its hexamer neighbors, which are contained within the alignment mask. Hence, after the initial subtomogram alignment, we selected only STA positions that were fully embedded in the lattice (meaning that they had six neighbors), in order to remove CA hexamers at the lattice edges from the analysis.

As the datasets from both systems substantially differed in size, we designed a strategy for using equally sized data subsets from each system. For System 1, we divided the full dataset (65,876 subvolumes after cleaning lattice edges) into three equal parts based on time of acquisition, and from each of these parts selected 20,000 subvolumes for alignments and generation of the final respective averages. The dataset acquired on System 2 contained 26,518 subvolumes, from which 20,000 were selected (for details see Materials and Methods and Supplementary Table 2). This approach yielded three and one equally-sized data-subsets for System 1 and System 2, respectively.

We then took advantage of the inherent local symmetries of the processed spherical EIAV CANC VLPs. We aligned the respective datasets using C6 symmetry. We then generated final averages for all subsets applying C1 symmetry (20,000 asymmetric units, a.u.), C2 symmetry (40,000 a.u.), C3 symmetry (60,000 a.u.) and C6 symmetry (120,000 a.u.), without any additional alignments. We estimated the resolutions for all generated structures using the 0.143 and 0.5 Fourier shell correlation (FSC) criterion and used the resulting values to make B-factor plots for both criteria (Figure 2). The B-factor plots showed mostly insignificant differences in resolution between the C1, C2, C3 and C6 symmetric averages of all data subsets from both systems (Figure 2 a and b). In seven out of eight measurements the mean resolution estimate for reconstructions obtained from System 1 was higher by 0.1-0.2 Å than that of System 2. For one measurement of the C2-symmetrized averages, the 0.5 FSC criterion showed a difference of 0.6 Å.

**Figure 2:**
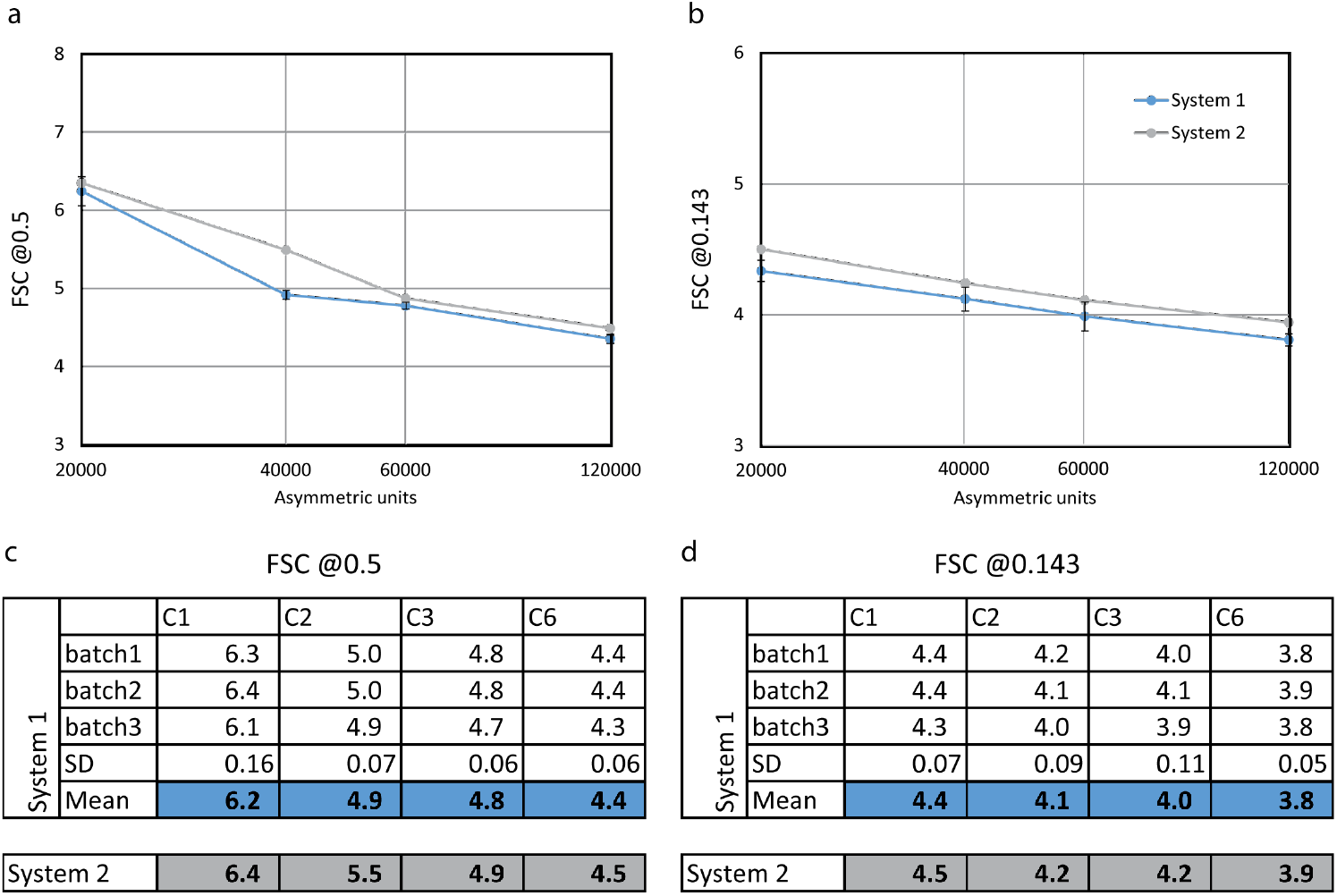
Subtomogram averaging resolution benchmark. Different symmetries (C1, C2, C3, C6) were applied to the maps for FSC measurements to assess the influence of dataset size and symmetry on resolution. **a)** FSC values at the 0.5 criterion plotted as a function of the number of asymmetric units. **b)** FSC values at 0.143 criterion plotted as a function of the number of asymmetric units. The curves for System 1 originate from the mean value of the measurements from the three subsets. X-axis in logarithmic scale in a,b. **c)** Resolution values in Å at the 0.5 FSC criterion, measured for different data-subsets from System 1 and System 2. **d)** Same as c, for values obtained at the 0.143 FSC criterion. SD is standard deviation.

However, upon inspection of the corresponding FSC curve from System 2 (Figure S2b, purple curve), we observed this larger difference to be caused by a local dip in FSC near the 0.5 criterion, rather than the FSC curve for this dataset being dramatically different from the others.

For the C6-symmetric averages the final resolution at the 0.143 FSC criterion was between 3.8 and 3.9 Å (Figure 2d, Figure S2). The three selected subsets of System 1 yielded resolutions of 3.8, 3.9, and 3.8 Å, while the dataset of System 2 resulted in a reconstruction at 3.9 Å. Importantly, upon inspection of the FSC curves for all data subsets, one of the System 1 subsets showed improved resolution compared to the other two subsets. (Figure S2). Hence, while the mean resolution obtained for System 1 was higher, the difference was less dramatic for two out of the three subsets from System 1.

Overall, the STA maps obtained for both systems were highly comparable in terms of quality. Differences in resolution of 0.1-0.2 Å can be considered rather negligible, especially in the context of a previous study, which benchmarked the influence of different tilt schemes on final map resolution (Turoňová et al., 2020). There, FSC results differed significantly more - by 0.3-2.8 Å - despite all data being acquired on the very same system. Hence, our results show that both of our systems perform comparably well at STA in the sub-4 Å resolution regime for small datasets, and that other experimental factors may dictate the achieved resolution. It is also important to point out that our results should not be interpreted as evidence that one system performs better than the other. Examples of experimental differences between the two datasets that may have affected the observed STA performance are described here. System 1 uses a cold FEG as electron source (E-CFEG), which provides a more coherent beam than the high-brightness Schottky FEG (XFEG) of System 2. The impact of a more coherent illumination on cryo-ET data acquisition has so far not been systematically explored. In case of the dataset acquired on System 2 the specimen condition might have been affected by increased deposition of contamination on the grid, due to multiple loading/ unloading events and shipping.

Overall, our evaluation allows one to assess the performance and suitability of the two systems for determining high-resolution cryo-ET and STA structures from moderately-sized datasets.

Next, we performed STA for the whole dataset acquired on System 1, using the AV3 subtomogram averaging routine. In addition to the particles fully embedded in the lattice, we also included particles at the edges, as long as they had at least 4 neighbors. By doing so, we aimed to maximize dataset size to determine the highest achievable resolution for the System 1 dataset using the AV3-based approach. We did not perform this step for System 2, as the data selected from System 2 for the benchmark presented in Figure 2 already contained 75% of all subtomograms, which would have meant repeating prior analysis with only a small increase in dataset size. For System 1, this approach yielded ~110,000 particles for further analysis. Crosscorrelation cleaning after the last alignment, to optimize the resolution determined via FSC, reduced the number of particles included in the final reconstruction to ~62,000 (~373,000 a.u). This resulted in a 3.4 Å resolution structure (Figure S3a). This corresponds to an increase of ~0.5 Å by using 3.1 times more data, compared to the resolutions shown for System 1 in Figure 2.

Since this structure was at the identical resolution as the previously solved highest-resolution structure of a retrovirus CA assembly using the AV3-based processing pipeline (EMPIAR-10164, processed in (Turoňová et al., 2017)), we reasoned that at this point this may represent the maximally achievable resolution given the data set size and the processing approach.

### Multiparticle refinement of System 1 and System 2 full datasets

To determine if employing an alternative processing approach could improve the final map resolution, we used a recently described software pipeline of Warp, Relion and Multiparticle refinement in M, which reported the highest resolution cryo-ET and STA structure of a retroviral CA assembly at 3.0 Å (EMPIAR-10164, processed in (Tegunov et al., 2021)). By doing so we aimed to assess the performance of System 1 in the sub-4 Å regime (and also System 2, although we predicted a lower resolution due to the significantly smaller overall dataset size).

M employs local tilt-series alignment and refinement of the CTF model, as well as utilization of multiple molecular species for this step. We therefore exploited the presence of tubular VLPs (consisting of C2-symmetric CA hexamers) within our tomograms (see also Figure 1a,b), which were distributed over the field of view in most tomograms and therefore represented valuable alignment features. Additionally, since the crosscorrelation cleaning approach was not applicable when using Relion, we attempted to increase the homogeneity of the subvolumes used for refinement by a more stringent exclusion of subvolumes that deviate from ideal lattice geometry. In order to retain only subvolumes containing the structurally most similar local CA hexamers we employed a cleaning strategy based on local geometry, similar to what has been previously used for mature HIV-1 and RSV CA assemblies (Mattei et al., 2016; Obr et al., 2021) (see also Figure S4 and Materials & Methods for more details).

For the dataset acquired on System 1, this processing pipeline resulted in a 2.9 Å resolution structure of the EIAV CA C6-symmetric hexamer from spherical VLPs (using ~466,000 a.u.) (Figure 3) and a 3.2 Å resolution reconstruction of the EIAV CA C2-symmetric hexamer from tubular VLPs (using ~107,000 a.u.) (Figure S3a-b), respectively.

**Figure 3:**
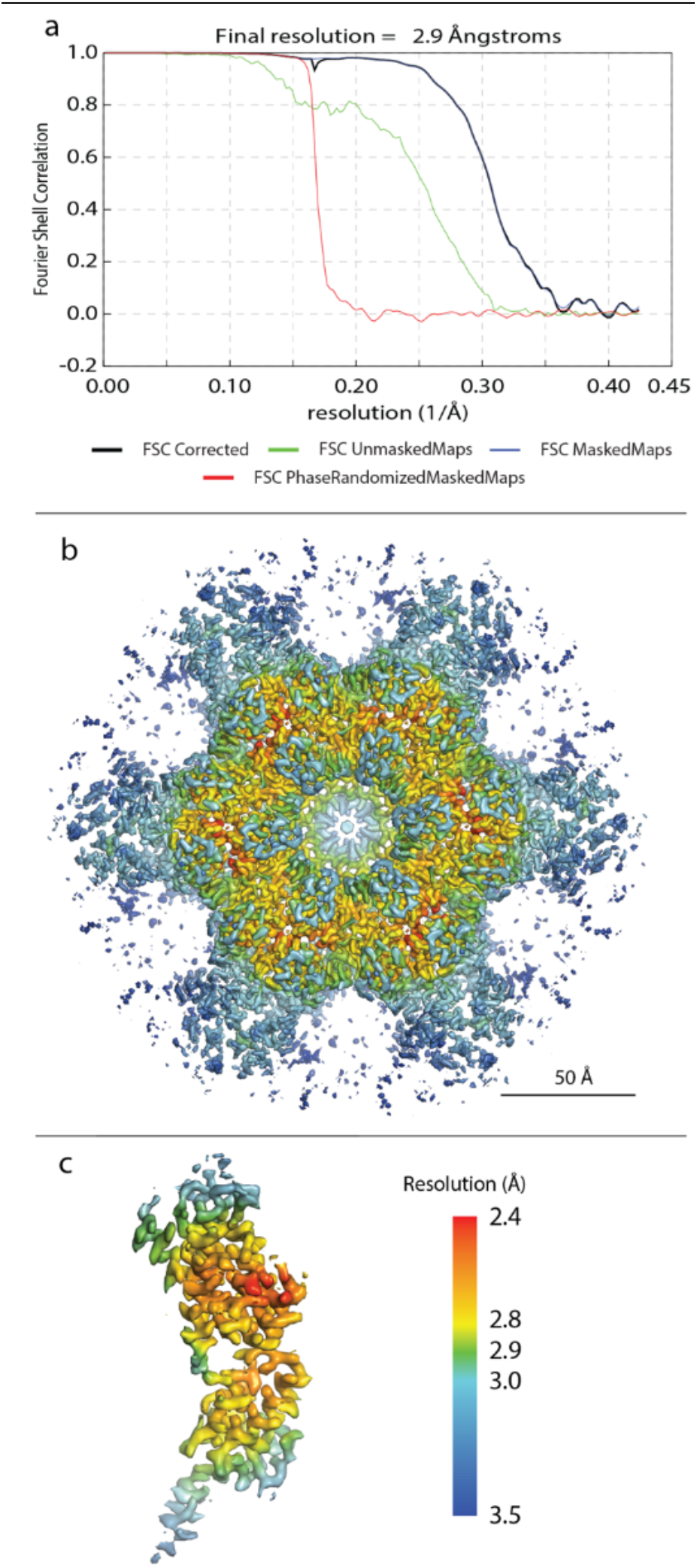
2.9 Å EIAV CA hexamer structure solved by multiparticle refinement using the full dataset acquired on System 1. **a)** FSC-curve. **b-c)** Structure of the full C6-symmetric hexamer (b) and one monomer (c) colored according to the local resolution as reported by M. The resolution range is depicted in the color bar.

While the structures from AV3-based and Relion/M-based processing cannot be directly compared due to a different cleaning approach, the increase in resolution is nevertheless notable and cannot alone be explained by the further increased subvolume quality, contributing to the final average. Our results demonstrate the impact of iterative local tilt-series alignment and CTF model refinement.

The local resolution map of the C6-symmetric hexamer reveals that the most stable regions of the hexameric lattice are located at the trimerization interface (Figure 3b,c). This is consistent with our previous work that identified the importance of this interface for immature EIAV lattice formation (Dick et al., 2020).

To verify the resolutions we obtained, and to test the effect of different masking strategies on FSC resolution estimates, we measured the resolution of our sub-3 Å map using two standard masks, as employed in other STA studies (Himes and Zhang, 2018; Schur et al., 2016; Tegunov et al., 2021; Turoňová et al., 2020, 2017). First, a cylindrical mask and second, a body-shaped mask (Figure S5a), which better matches the shape of a single CA hexamer. For both masks, the FSC curves were nearly identical (Figure S5b) and importantly, led to the same resolution estimates. This suggests that use of either masking approach, as long as the mask encompasses the whole central unit, and includes sufficient Gaussian smoothing, is acceptable for resolution estimates of continuous retroviral lattices.

We also subjected the dataset from System 2 to the same Relion/M processing pipeline. This dataset contained only 45% and 35% of the number of subvolumes from spherical and tubular VLPs respectively compared to data from System 1. The workflow employing multiparticle refinement resulted in a 3.3 Å structure of the EIAV CA C6 symmetric hexamer from spherical VLPs (~195,000 a.u.) (Figure S3b) and a 3.6 Å structure of the EIAV CA C2 symmetric hexamer from tubular VLPs (from ~37,000 a.u.) (Figure S3d).

Comparison of the C6-symmetric structures determined in this study, to the previously solved structure from an identical sample from data acquired on a Titan Krios G1 equipped with a Bioquantum K2 (Dick et al., 2020), revealed visible, but still very subtle differences (Figure 4), despite the difference in resolution of up to 0.8 Å. An important consideration in this comparison is that the dataset acquired on the Gatan K2 system was not processed via multiparticle refinement, but instead using the pipeline shown in Figure 2 and Figure S3a.

**Figure 4.**
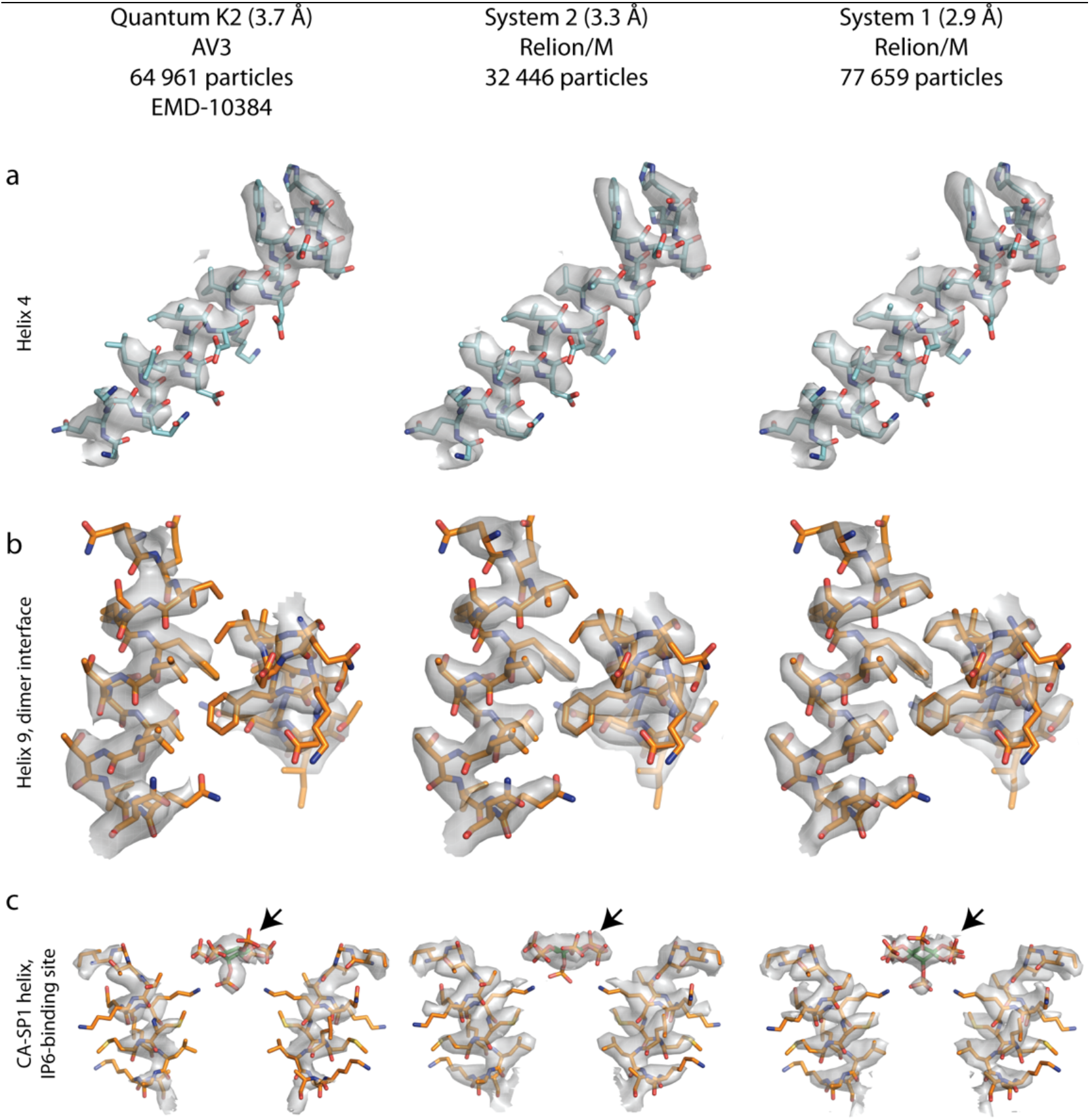
Gallery of EIAV-CANC structures solved using different detectors. Representative EM-densities from the structures generated via Relion/M in this study (center and right) and a structure determined previously using novaCTF/AV3 and the previous generation of the Gatan DED (left). The model for EIAV CA (pdb 6T64) fitted into the EM densities is consistent with all three structures. Notably, the density for IP6, a small molecule employed by EIAV for stabilizing the immature CA lattice, is best resolved within the lowest-resolution structure. **a)** Helix 4 in the N-terminal domain of EIAV CA; **b)** helix 9 in the C-terminal domain of EIAV CA, forming an important dimeric inter-hexamer interface; **c)** Inositolhexakiphosphate (IP6) binding pocket, for clarity only two opposite helices from the 6-helix bundle are shown. IP6 is annotated by an arrow.

## Conclusions

Here we describe our assessment of the potential of two different EFTEM setups combining narrow-band energy filtering and the newest generation of DEDs for high-resolution structure determination using cryo-ET and STA.

Our results clearly show that both systems support high-resolution cryo-ET data collection and perform comparably in the ~4-6 Å resolution regime for our STA benchmark. Furthermore, our reconstruction at 2.9 Å demonstrates that System 1 is capable of collecting cryo-ET data that can be processed to sub-3 Å using STA. While the dataset size for System 2 did not allow us to achieve the same resolution, the sub-3 Å subtomogram averaging capabilities of the K3 have recently been shown in a different publication (Tegunov et al., 2021).

However, the number of specimens which can currently reach sub-3Å resolution via cryo-ET and STA is likely constrained to *in vitro* (*biochemically pure*) samples. Considering the increased use of cryo-ET for *in situ* structure determination, we believe that our observations at the resolutions between 4-6 Å, using smaller datasets, are of higher relevance. Based on the results presented here and in previous studies, appropriate cryo-ET data acquisition conditions, e.g. choice of the tilt scheme (Turoňová et al., 2020), might represent a more dominant factor for determining cryo-ET data quality, than the choice between systems equipped with latest generation

DEDs. Since pipelines supporting native EER processing were not available at the time of this publication, we have not explored all the options that the new data format makes available. Future studies should help determine if further advantages of the EER format are beneficial to STA.

## Acknowledgements

This work was funded by the Austrian Science Fund (FWF) grant P31445 to F.K.M.S and the National Institute of Allergy and Infectious Diseases under awards R01AI147890 to R.A.D. This research was also supported by the Scientific Service Units (SSUs) of IST Austria through resources provided by Scientific Computing (SciComp), the Life Science Facility (LSF), and the Electron Microscopy Facility (EMF). We thank Dustin Morado for providing the software SubTOM for data processing.

## Data availability

The electron microscopy density maps and representative tomograms have been deposited in the Electron Microscopy Data Bank (accession codes EMD-14031, EMD-14032, EMD-14033, EMD-14034, EMD-14035, EMD-14036, and EMD-14037). The EER movies and tilt series acquired on System 1 have been deposited to the Electron Microscopy Public Image Archive under the accession code: EMPIAR-10889.

## Materials & Methods

### In vitro assembly of EIAV VLPs

EIAV CANC VLPs were assembled as previously described (Dick et al., 2020). Briefly, 30 μL assembly reactions containing 50 μM purified CASPNC protein, 500 mM NaCl, 10 μM IP6, 2 mM TCEP, and 10 μM GT50 oligonucleotide were dialyzed against buffer containing 20 mM MES pH 6.2, 100 mM NaCl, 2 mM TCEP, and 10 μM IP6 for 4 hours at 4°C. To assess assembly, 5 μL of the assembly reaction was spotted on glow discharged (15 mA, 30 sec) grids (formvar/carbon 200), followed by 2% uranyl acetated staining, and imaging on a Morgagni transmission electron microscope.

### Sample preparation for cryo-ET

Cryo-EM grids with CASPNC VLPs were prepared using a Leica GP2 plunger. 2.5 μl of EIAV CASPNC VLPs, mixed with 10nm colloidal gold, were applied to glow discharged 2/2-C C-flat grids immediately before vitrification. The grids were then back-side blotted for 3.5 s at 10°C and ~95% humidity, subsequently plunged into liquid ethane, and then transferred to liquid nitrogen for storage. System 1 and System 2 data acquisition was performed on the same grid.

### Krios G4 Selectris X Falcon 4 (System 1) data acquisition

The first dataset was acquired using a Krios G4 with cold field emission gun (CFEG; Thermo Fisher Scientific) and a Selectris X energy filter with a Falcon 4 detector (Thermo Fisher Scientific). Data were collected with TEM Tomography 5 software (Thermo Fisher Scientific). Tilt-series were recorded at a nominal magnification of ×105,000, corresponding to a pixel size of 1.179 Å. A dose-symmetric scheme was used to collect tilt-series from −60° to 60° at a step size of 3°. The CFEG was automatically flashed every ~8 h. Data were collected using the EER format of Falcon 4. The total dose per tilt was 3.5e/ Å^2^. A 10eV slit was used. Eucentric height was performed once for the entire grid square using the stage tilt method in TEM Tomography 5 software. Regions of interest for data collection were determined manually. Tracking and focusing was applied before and after acquisition of each tilt step. The energy filter zero-loss peak was tuned once prior to data acquisition. The target defocus for each tilt series was changed over a range of −0.75 to −3.25 um in steps of 0.25 um.

### Krios G3i Bioquantum K3 (System 2) data acquisition

The second dataset was acquired using a Krios G3i with a XFEG electron source (Thermo Fisher Scientific) and a BioQuantum energy filter with a K3 detector (Gatan). The grid was mapped at the lowest possible magnification, suitable grid squares were identified and mapped at the lowest SA magnification, omitting the grid squares already collected on by System 1. Tilt series were acquired using the Tilt Controller in SerialEM (Mastronarde, 2005). The calibrated pixel size was 0.822 Ångstrom, and the exposure time 0.096 seconds, which allowed for 7 frames to be saved in LZW compressed uncorrected TIF format. A dose-symmetric scheme was used to collect tilt-series from −60° to 60° at a step size of 3°. The total dose per tilt was 3.7 e/Å^2^. A 20eV slit was used. For each tilt series the SerialEM Eucentric Fine procedure was run to set stage height, followed by alignment of the point of interest. Tracking and focusing was applied for each tilt step, at the end of each tilt series the energy filter zero-loss peak was refined and a new K3 hardware dark reference acquired. The target defocus for each tilt series was changed over a range of −0.75 to −3.25 um in steps of 0.25 um.

### NovaCTF/AV3 processing

The image processing workflow is schematically depicted in Figure S1. The initial processing for both System 1 and System 2 datasets was done identically, with the exception that the raw EER files obtained via System 1 were aligned and summed into mrc stacks using Relion 3.1. The tif frames generated by System 2 were aligned and summed using the alignframes plugin in IMOD.

Prior to further processing, bad tilts (e.g. images that shifted significantly during acquisition or due to a blocked beam at high tilts) were removed. The stacks were then low-pass filtered according to the respective exposure dose in individual tilts (Grant and Grigorieff, 2015). IMOD (Kremer et al., 1996) was used for tilt series alignment and to generate 8x binned tomograms, filtered using the SIRT-like filter in IMOD for manual picking. Defocus was estimated using CTFFIND4 (Rohou and Grigorieff, 2015). The full tomograms were reconstructed in NovaCTF (Turoňová et al., 2017) with simultaneous 3D CTF correction using the multiplication algorithm. The slab thickness was set to 15 nm.

The initial positions for subtomogram averaging were obtained as previously described (Dick et al., 2020). Briefly, the centers of VLPs were marked and saved as models in 3dmod. Subtomograms were then extracted at positions on the surface of a sphere with a radius corresponding to the dimension of the respective VLP, using a custom MATLAB script.

Subtomogram averaging and alignment was performed in the AV3/TOM-based package SubTOM (https://github.com/DustinMorado/subTOM), as described (Dick et al., 2020). Throughout processing we also employed Dynamo functionalities (Castano-Diez et al., 2012). Specifically, subtomogram alignment was done at bin8, bin4, bin2 and bin1, while gradually advancing the low-pass filter and decreasing the Euler angle scanning step and range. The initial reference was created de novo using a single VLP. The initial reference was then symmetrized (C6) and used to align subtomograms from all VLPs of a single high-defocus tomogram. The resulting reference was used as a starting template for the alignment of the whole dataset at bin8. Following alignments, subvolumes with overlapping positions, positions which diverged away from the VLP surface, as well as positions with low cross-correlation were removed.

As described in the results, for the analysis of the data subsets, only subtomograms, which were not located at the edges of the lattice, were selected. Specifically, we removed subtomograms that had fewer than 6 neighbors after alignment at bin8 (Figure S4). A conservative lowpass filter of 1/30 Å or lower was used up to the last iteration of bin4 processing to avoid over-refinement. After bin4 alignment subtomograms with grey scale values differing by more than 2 sigma from the mean of the entire dataset, were removed.

To generate equally sized datasets, the cleaned data obtained on System 1 was split into three equal parts based on time of acquisition, and 21,000 subtomograms were randomly selected from each dataset for further processing. For the dataset obtained on System 2, only one subset with 21,000 random subtomograms was generated due to the limiting amount of collected data. The four subsets were split into their respective odd and even half sets, which were then processed independently at bin2 with two rounds of alignments. After bin2 alignments 20,000 particles with the best cross-correlation values for each subset were retained. Finally, 3 rounds of alignment were performed at bin1.

To keep the processing parameters consistent between datasets from both systems, and since the expected resolution was far from the physical Nyquist frequency in the System 2 dataset (0.822 Å /pix corresponds to the Nyquist frequency of 1/1.644 Å), the subtomograms were cropped in Fourier space to approximately match the pixel size at which the Falcon 4 dataset was acquired (1.179 Å).

To obtain relevant subtomogram alignment parameters from the tubular VLPs for subsequent refinement in Relion and M, the processing in subTOM for tubes was done analogically to the processing for spheres, except using C2-symmetry and stopping after bin4 alignment, before proceeding with multiparticle refinement.

For processing the full System 2 dataset we included all subtomograms that had at least 4 neighbors after alignment at bin8. In addition, after the final alignment, the particles were cleaned using a crosscorrelation threshold to maximize the resolution at the 0.143 FSC criterion.

### Multiparticle refinement

For the System 1 dataset acquired on the Falcon 4 DED, EER files were summed using Relion 3.1 to yield 5 TIF frames for each tilt. Computed (Falcon 4) and raw (K3) frames were processed using Warp. The tilt series alignments from Etomo and subtomogram alignments from the NovaCTF/AV3 pipeline were imported into Warp version nb20201104 using the dynamo2m script package (https://github.com/alisterburt/dynamo2m), and used for subtomogram reconstruction. Subtomograms were reconstructed at bin4 using Warp, and Autorefine 3D was performed in Relion 3.0.7e. These steps were repeated at bin2 to yield the starting positions for multi-particle refinement in M. For the multiparticle refinement we excluded particles, which were outliers in terms of local lattice geometry. Briefly, we excluded hexamers which had less than 3 neighbors fulfilling the geometry restraints specified in Figure S4b,c. Subsequently, five rounds of multiparticle refinement were performed, gradually extending the refinement options (Supplementary Table 3, Supplementary Table 4). As previously described, to use similarly sized boxes and masks for both systems, we processed the data from System 1 at its nominal pixel size (1.179 Å) and the data from System 2 at a pixel size of 1.1508 Å (corresponding to a cropping factor of 1.4). The refinement was performed analogously for both datasets, with the only exception being the different setup of tiles for image and volume warp, to account for the rectangular shape of the K3 camera.

### Resolution estimation

The resolution of the final averages obtained via the NovaCTF/AV3 pipeline was determined by a phase-randomized mask-corrected FSC using a Gaussian filtered cylindrical mask (for example see Figure S5a).

To verify the resolution of the map obtained using Multiparticle refinement, we assessed two different masks - a body mask generated using the pdb model of EIAV CA hexamer (pdb 6T64) (Figure S5a), and the same tight cylindrical mask around one hexamer, which was used for the FSC of the AV3 pipeline results (Figure S5a). The resolution was measured in MATLAB, as well as Relion. The results were consistent and supportive of the reported resolution regardless of which mask or software was used for the FSC calculation (Figure S5b-c). The half maps of the M refinement were then combined, B-factor sharpened and filtered to the measured resolution in M using the cylindrical mask (Figure S5a, left) (Rosenthal and Henderson, 2003). Visualization of tilt series, tomograms and EM-densities was performed in MATLAB, IMOD (Kremer et al., 1996), UCSF Chimera (Pettersen et al., 2004), and Pymol (Schrodinger LLC, 2010).

## Supplementary Figures

**Figure S1:**
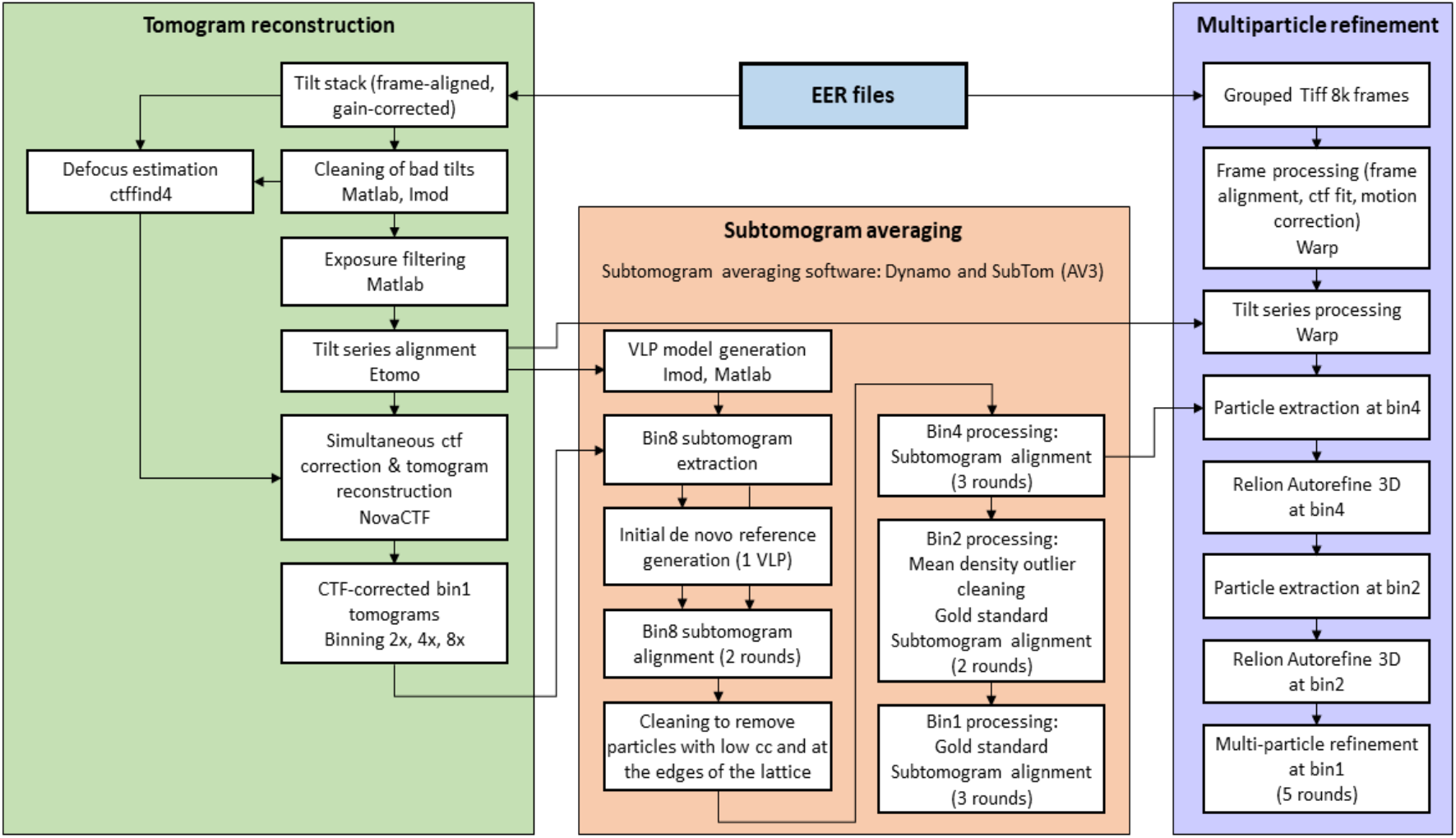
Data processing chart Diagram of the image processing workflows employed in this study. The green sector indicates pre-processing and tomogram reconstruction, the orange sector contains subtomogram averaging performed in MATLAB, Dynamo and subTOM (AV3), and the purple sector contains all steps performed in Relion and M.

**Figure S2.**
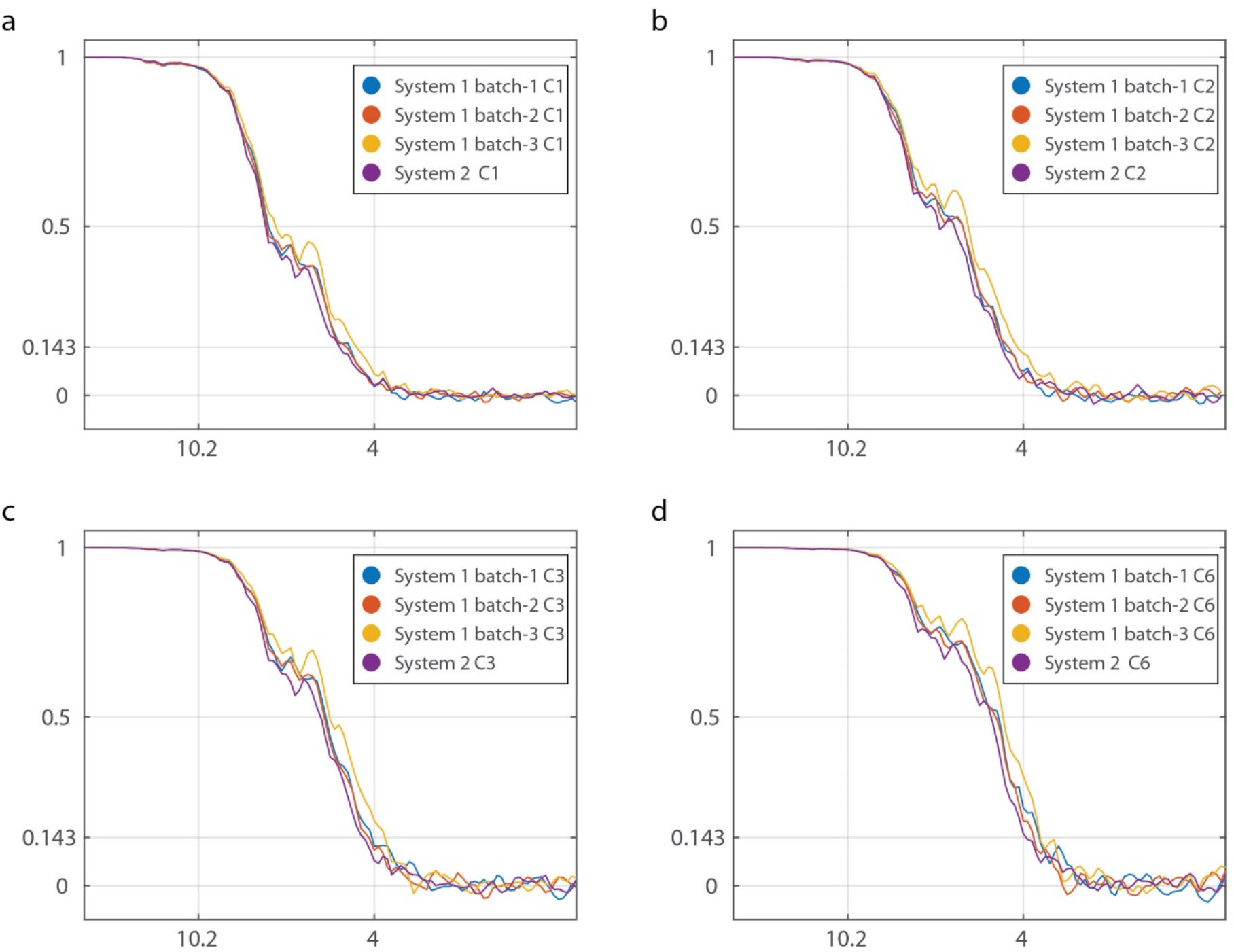
Fourier shell correlation of symmetrized maps solved using System 1 and System 2, corresponding to data shown in Figure 2. **a-d** FSCs of symmetrized maps **a** – C1, **b** – C2, **c** – C3, and **d** – C6. The FSC crossings at 0.5 and 0.143 correspond to Figure 2 and the values in Supplementary Table 2.

**Figure S3:**
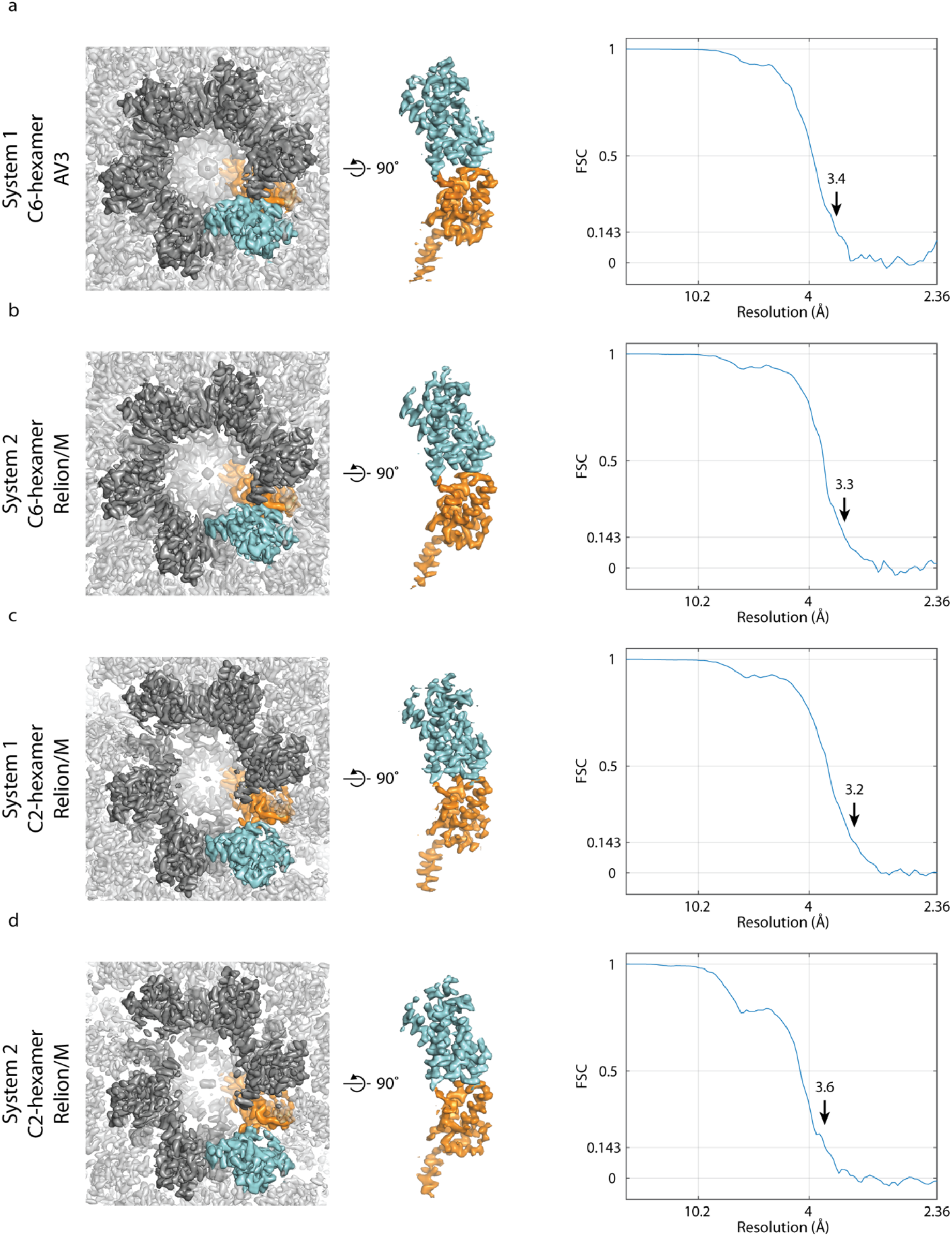
Cryo-ET maps solved by STA using data from System 1 and System 2. **a-d)** Isosurface representations of sharpened cryo-ET maps, generated either via the NovaCTF/AV3-based or Relion/M-based workflows. The origin of the data (System 1 or 2), the respective average (C6-hexamer from spherical VLPs, or C2-hexamer from tubular VLPs) and the used software are shown at the left. The central CA hexamer of the solved structures is shown in dark gray, with one CA monomer within the hexamer highlighted with cyan and orange colors for the NTD and CTD, respectively. The corresponding FSC-curves are shown on the right. **a)** EIAV C6-symmetric CA hexamer solved by the NovaCTF/AV3 pipeline from the full System 1 dataset. The obtained resolution at the 0.143 FSC criterion is 3.4Å. **b)** EIAV C6-symmetric CA hexamer solved by Multiparticle refinement from the full System 2 dataset. The obtained resolution at the 0.143 FSC criterion is 3.3Å. **c)** EIAV C2-symmetric CA hexamer solved by Multiparticle refinement from the full System 1 dataset. The obtained resolution at the 0.143 FSC criterion is 3.2Å. **d)** EIAV C2-symmetric CA hexamer solved by Multiparticle refinement from the full System 2 dataset. The obtained resolution at the 0.143 FSC criterion is 3.6Å.

**Figure S4:**
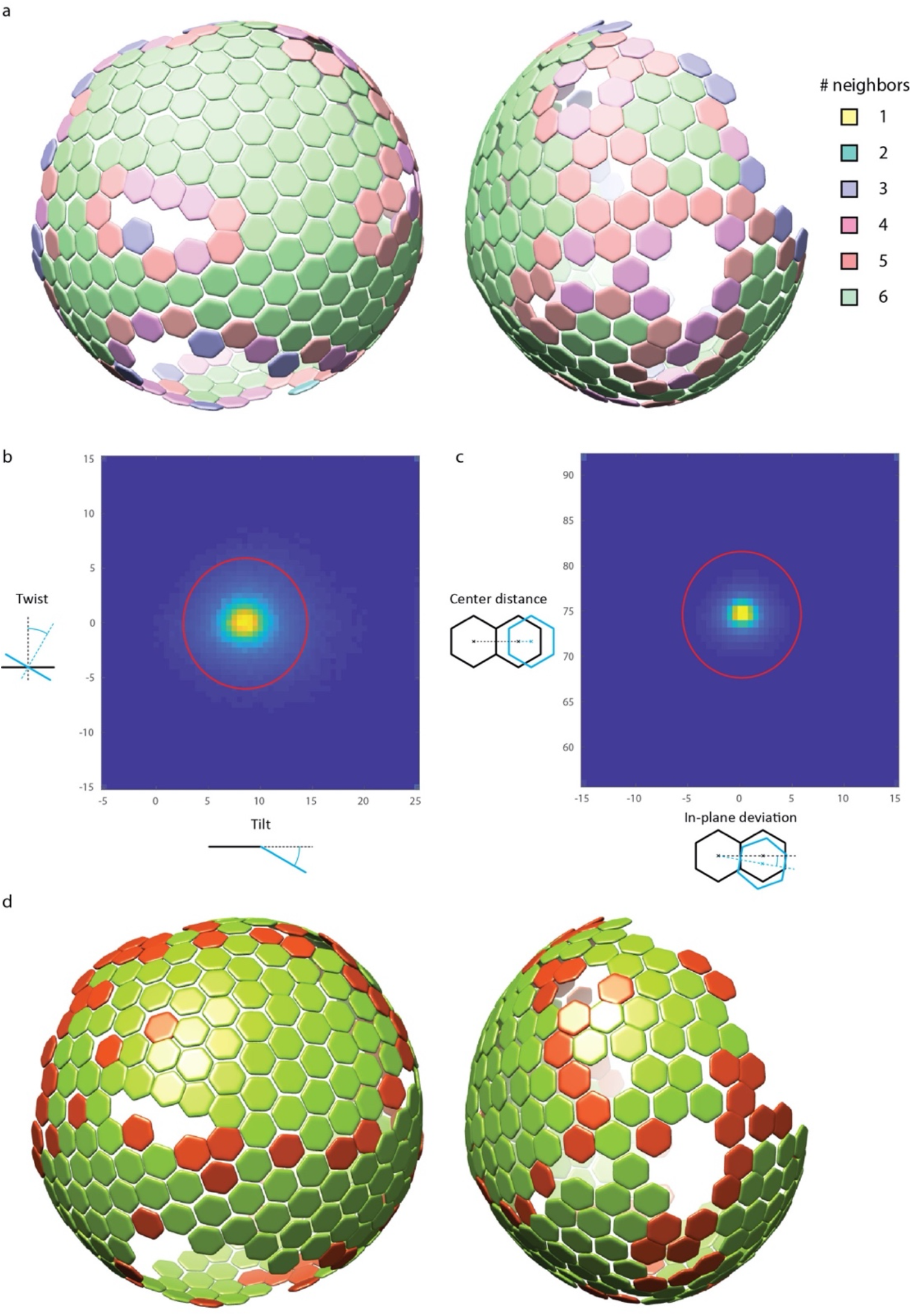
Geometry cleaning of the EIAV immature CANC lattice. **a)** Schematic representation of hexamer positions for a given VLP. Color code corresponds to the number of neighboring hexamers: green - 6; salmon - 5; pink – 4; purple – 3; light blue – 2; yellow – 1. **b)** A plot visualizing the tilt and twist angles for the individual hexamer-hexamer neighbors. The red circle shows the threshold used for cleaning. **c)** A plot visualizing the inplane angle deviation and the center-to-center distance for the individual hexamer-hexamer neighbors. The red circle shows the threshold used for the cleaning. **d**) Same as **a**, shown in green positions that fulfill conditions visualized in **b** and **c** for at least 3 neighbors; the remaining outlier positions are shown in red.

**Figure S5:**
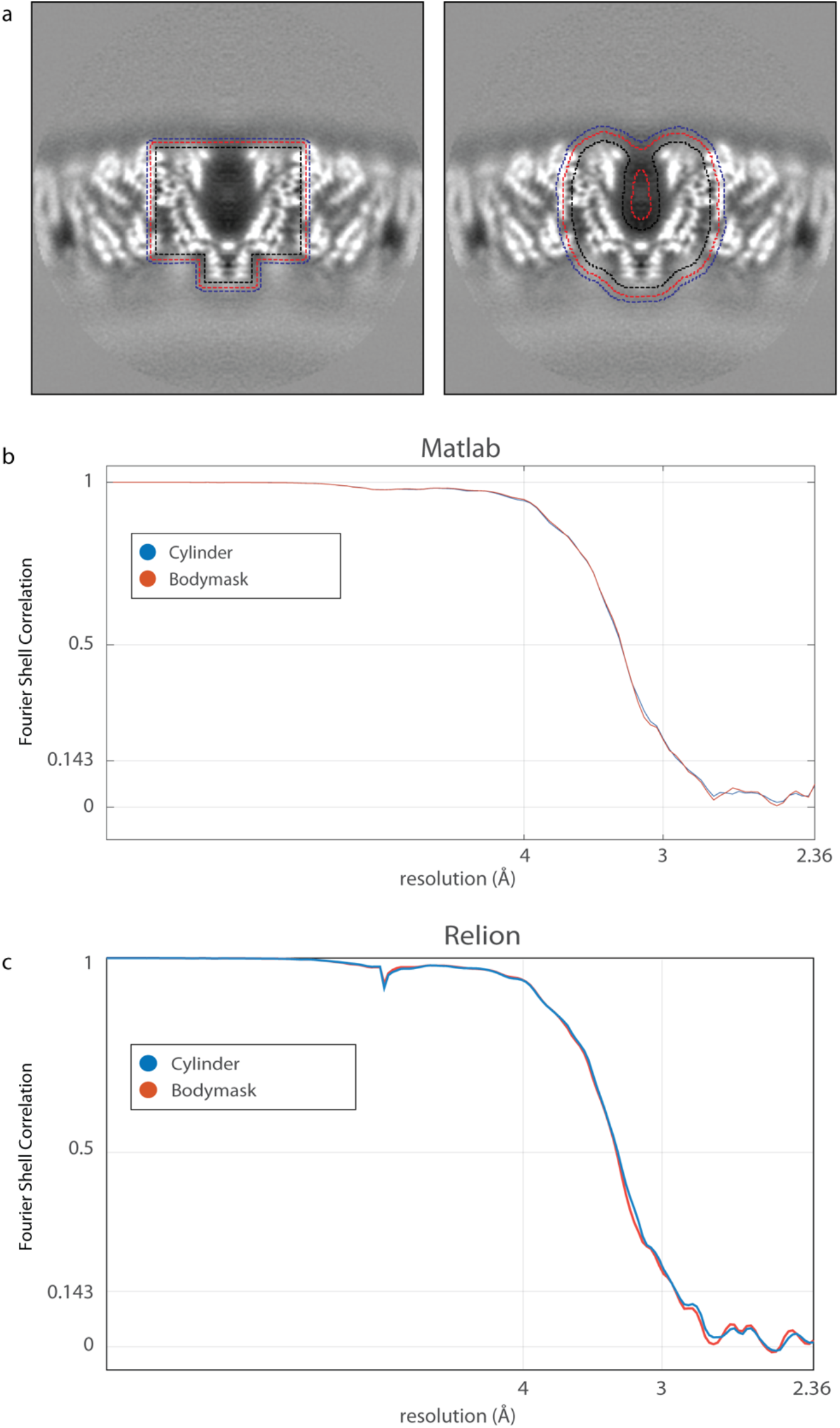
Effects of the mask on the FSC. **a)** Central slice through the C6-symmetric map obtained by Multiparticle refinement for System 1. The boundary of the masks, either cylindrical or body-shaped is shown by dotted lines. Black dotted line –isosurface threshold at 1.0, red dotted line – isosurface threshold at 0.67, and blue dotted line – isosurface threshold at 0.33. **b)** Phase-randomized and masked FSC performed using MATLAB. **c** Phase-randomized, masked FSC in Relion 3.0.7.

## References

Bäuerlein, F.J.B., Baumeister, W., 2021. Towards Visual Proteomics at High Resolution. J. Mol. Biol. 433, 167187. https://doi.org/10.1016/j.jmb.2021.167187

Bharat, T.A.M., Russo, C.J., Löwe, J., Passmore, L.A., Scheres, S.H.W., 2015. Advances in Single-Particle Electron Cryomicroscopy Structure Determination applied to Sub-tomogram Averaging. Structure 23, 1743–1753. https://doi.org/10.1016/j.str.2015.06.026

Bouvette, J., Liu, H.F., Du, X., Zhou, Y., Sikkema, A.P., da Fonseca Rezende e Mello, J., Klemm, B.P., Huang, R., Schaaper, R.M., Borgnia, M.J., Bartesaghi, A., 2021. Beam image-shift accelerated data acquisition for near-atomic resolution single-particle cryo-electron tomography. Nat. Commun. 12, 1–11. https://doi.org/10.1038/s41467-021-22251-8

Bush, D.L., Vogt, V.M., 2014. In Vitro Assembly of Retroviruses. Annu. Rev. Virol. 1, 561–580. https://doi.org/doi:10.1146/annurev-virology-031413-085427

Castano-Diez, D., Kudryashev, M., Arheit, M., Stahlberg, H., 2012. Dynamo: a flexible, user-friendly development tool for subtomogram averaging of cryo-EM data in high-performance computing environments. J Struct Biol 178, 139–151. https://doi.org/10.1016/j.jsb.2011.12.017

Chen, M., Bell, J.M., Shi, X., Sun, S.Y., Wang, Z., Ludtke, S.J., 2019. A complete data processing workflow for cryo-ET and subtomogram averaging. Nat. Methods 16, 1161–1168. https://doi.org/10.1038/s41592-019-0591-8

Chreifi, G., Chen, S., Metskas, L.A., Kaplan, M., Jensen, G.J., 2019. Rapid tilt-series acquisition for electron cryotomography. J. Struct. Biol. 205, 163–169. https://doi.org/10.1016/J.JSB.2018.12.008

Dick, R.A., Xu, C., Morado, D.R., Kravchuk, V., Ricana, C.L., Lyddon, T.D., Broad, A.M., Feathers, J.R., Johnson, M.C., Vogt, V.M., Perilla, J.R., Briggs, J.A.G., Schur, F.K.M., 2020. Structures of immature EIAV Gag lattices reveal a conserved role for IP6 in lentivirus assembly. PLoS Pathog. 16. https://doi.org/10.1371/journal.ppat.1008277

Eisenstein, F., Danev, R., Pilhofer, M., 2019. Improved applicability and robustness of fast cryoelectron tomography data acquisition. J. Struct. Biol. 208, 107–114. https://doi.org/10.1016/j.jsb.2019.08.006

Förster, F., Medalia, O., Zauberman, N., Baumeister, W., Fass, D., 2005. Retrovirus envelope protein complex structure in situ studied by cryo-electron tomography. Proc Natl Acad Sci U S A 102, 4729–4734. https://doi.org/10.1073/pnas.0409178102

Grant, T., Grigorieff, N., 2015. Measuring the optimal exposure for single particle cryo-EM using a 2.6 Å reconstruction of rotavirus VP6. Elife 4, e06980. https://doi.org/10.7554/eLife.06980

Hagen, W.J.H., Wan, W., Briggs, J.A.G., 2017. Implementation of a cryo-electron tomography tilt-scheme optimized for high resolution subtomogram averaging. J. Struct. Biol. 197, 191–198. https://doi.org/10.1016/j.jsb.2016.06.007

Himes, B.A., Zhang, P., 2018. emClarity: software for high-resolution cryo-electron tomography and subtomogram averaging. Nat. Methods 15, 955–961. https://doi.org/10.1038/s41592-018-0167-z

Kremer, J.R., Mastronarde, D.N., McIntosh, J.R., 1996. Computer Visualization of Three-Dimensional Image Data Using IMOD. J Struct Biol 116, 71–76. http://dx.doi.org/10.1006/jsbi.1996.0013

Mastronarde, D.N., 2005. Automated electron microscope tomography using robust prediction of specimen movements. J. Struct. Biol. 152, 36–51. https://doi.org/10.1016/j.jsb.2005.07.007

Mattei, S., Glass, B., Hagen, W.J.H., Kräusslich, H.G., Briggs, J.A.G., 2016. The structure and flexibility of conical HIV-1 capsids determined within intact virions. Science (80-.). 354, 1434–1437. https://doi.org/10.1126/science.aah4972

Nakane, T., Kotecha, A., Sente, A., McMullan, G., Masiulis, S., Brown, P.M.G.E., Grigoras, I.T., Malinauskaite, L., Malinauskas, T., Miehling, J., Uchański, T., Yu, L., Karia, D., Pechnikova, E. V, de Jong, E., Keizer, J., Bischoff, M., McCormack, J., Tiemeijer, P., Hardwick, S.W., Chirgadze, D.Y., Murshudov, G., Aricescu, A.R., Scheres, S.H.W., 2020. Single-particle cryo-EM at atomic resolution. Nature 587, 152–156. https://doi.org/10.1038/s41586-020-2829-0

O’Reilly, F.J., Xue, L., Graziadei, A., Sinn, L., Lenz, S., Tegunov, D., Blötz, C., Singh, N., Hagen, W.J.H., Cramer, P., Stülke, J., Mahamid, J., Rappsilber, J., 2020. In-cell architecture of an actively transcribing-translating expressome. Science (80-.). 369, 554–557. https://doi.org/10.1126/science.abb3758

Obr, M., Ricana, C.L., Nikulin, N., Feathers, J.-P.R., Klanschnig, M., Thader, A., Johnson, M.C., Vogt, V.M., Schur, F.K.M., Dick, R.A., 2021. Structure of the mature Rous sarcoma virus lattice reveals a role for IP6 in the formation of the capsid hexamer. Nat. Commun. 12, 3226. https://doi.org/10.1038/s41467-021-23506-0

Obr, M., Schur, F.K.M., 2019. Structural analysis of pleomorphic and asymmetric viruses using cryo-electron tomography and subtomogram averaging, in: Advances in Virus Research. pp. 117–159. https://doi.org/10.1016/bs.aivir.2019.07.008

Pettersen, E.F., Goddard, T.D., Huang, C.C., Couch, G.S., Greenblatt, D.M., Meng, E.C., Ferrin, T.E., 2004. UCSF Chimera--a visualization system for exploratory research and analysis. J Comput Chem 25, 1605–1612. https://doi.org/10.1002/jcc.20084

Rohou, A., Grigorieff, N., 2015. CTFFIND4: Fast and accurate defocus estimation from electron micrographs. J Struct Biol 192, 216–221. http://dx.doi.org/10.1016/j.jsb.2015.08.008

Rosenthal, P.B., Henderson, R., 2003. Optimal Determination of Particle Orientation, Absolute Hand, and Contrast Loss in Single-particle Electron Cryomicroscopy. J Mol Biol 333, 721–745. https://doi.org/10.1016/j.jmb.2003.07.013

Sanchez, R.M., Zhang, Y., Chen, W., Dietrich, L., Kudryashev, M., 2020. Subnanometer-resolution structure determination in situ by hybrid subtomogram averaging - single particle cryo-EM. Nat. Commun. 11, 3709. https://doi.org/10.1038/s41467-020-17466-0

Scaramuzza, S., Castaño-Díez, D., 2021. Step-by-step guide to efficient subtomogram averaging of virus-like particles with Dynamo. PLOS Biol. 19, e3001318.

Schrodinger LLC, 2010. The PyMOL Molecular Graphics System, Version 1.3r1.

Schur, F.K., 2019. Toward high-resolution in situ structural biology with cryo-electron tomography and subtomogram averaging. Curr. Opin. Struct. Biol. 58, 1–9. https://doi.org/10.1016/j.sbi.2019.03.018

Schur, F.K.M., Obr, M., Hagen, W.J.H., Wan, W., Jakobi, A.J., Kirkpatrick, J.M., Sachse, C., Kräusslich, H.G., Briggs, J.A.G., 2016. An atomic model of HIV-1 capsid-SP1 reveals structures regulating assembly and maturation. Science (80-.). 353, 506–508. https://doi.org/10.1126/science.aaf9620

Tan, A., Pak, A.J., Morado, D.R., Voth, G.A., Briggs, J.A.G., 2021. Immature HIV-1 assembles from Gag dimers leaving partial hexamers at lattice edges as potential substrates for proteolytic maturation. Proc. Natl. Acad. Sci. 118, e2020054118. https://doi.org/10.1073/pnas.2020054118

Tegunov, D., Xue, L., Dienemann, C., Cramer, P., Mahamid, J., 2021. Multi-particle cryo-EM refinement with M visualizes ribosome-antibiotic complex at 3.5 Å in cells. Nat. Methods 18, 186–193. https://doi.org/10.1038/s41592-020-01054-7

Turoňová, B., Hagen, W.J.H., Obr, M., Mosalaganti, S., Beugelink, J.W., Zimmerli, C.E., Kräusslich, H.-G., Beck, M., 2020. Benchmarking tomographic acquisition schemes for high-resolution structural biology. Nat. Commun. 11, 876. https://doi.org/10.1038/s41467-020-14535-2

Turoňová, B., Schur, F.K.M., Wan, W., Briggs, J.A.G., 2017. Efficient 3D-CTF correction for cryoelectron tomography using NovaCTF improves subtomogram averaging resolution to 3.4 Å. J. Struct. Biol. 199, 187–195. https://doi.org/10.1016/j.jsb.2017.07.007

Wan, W., Briggs, J.A.G., 2016. Chapter Thirteen - Cryo-Electron Tomography and Subtomogram Averaging, in: Crowther, R.A. (Ed.), Methods in Enzymology. Academic Press, pp. 329–367. http://dx.doi.org/10.1016/bs.mie.2016.04.014

Yip, K.M., Fischer, N., Paknia, E., Chari, A., Stark, H., 2020. Atomic-resolution protein structure determination by cryo-EM. Nature 587, 157–161. https://doi.org/10.1038/s41586-020-2833-4

Zhang, K., Pintilie, G.D., Li, S., Schmid, M.F., Chiu, W., 2020. Resolving individual atoms of protein complex by cryo-electron microscopy. Cell Res. 30, 1136–1139. https://doi.org/10.1038/s41422-020-00432-2

